# Mathematical modelling of megakaryopoiesis in Mpl-deficient and continuously thrombopoietin-stimulated mice points to an unknown control mechanism

**DOI:** 10.1101/2020.05.29.123489

**Authors:** Hans H. Diebner, Andrea Gottschalk, Christoph Baldow, Markus Klose, Ingmar Glauche

## Abstract

Thrombopoietin (TPO) is the ligand of the Mpl receptor and the key regulator of megakaryopoiesis and platelet production. A loss or gain of the TPO-receptor function affects haematopoiesis and results in severe diseases in humans. Appropriate mouse strains are available to mimic both myeloproliferative neoplasm (MPN) and congenital amegakaryocytic thrombocytopenia (CAMT) resulting from TPO overexpression or knockout of TPO receptor Mpl on megakaryocytes and platelets, respectively. However, at a quantitative level it is not understood, how the known regulations can establish the impaired but stable disease phenotypes.

Starting out from an established mathematical model for megakaryopoiesis, we aim to adapt it to both the healthy situation and to distinct diseased phenotypes mimicking MPN. We thereby identify, that some of the model parameters are invariant with respect to the mouse strain while others have to be estimated in a strain-dependent manner. A systematic process of parameter identification provides strong evidence that the well-known excess production of megakaryocytes and early progenitors in MPN is either directly contingent on Mpl expression of platelets and megakaryocytes or, alternatively, that the knockout of Mpl is not as precisely restricted to megakaryocytes and platelets but may also effect their progenitors.

In conclusion, our analysis hints towards an opaque control mechanism rendering megakaryopoiesis at a yet unknown level, and awaiting further experimental evaluation.

## Introduction

The cytokine thrombopoietin (TPO) plays an essential role in megakaryopoiesis and thrombocyte (platelet) production^1,2^. TPO is mainly produced constitutively by the liver but is also inducible during inflammation^3^. By binding to its receptor, i.e. the cellular homologue of the myeloproliferative leukaemia virus oncogene (Mpl), TPO controls the homeostatic levels of primitive haematopoietic cells as well as myeloid and erythroid progenitors in the bone marrow. Most importantly, TPO also regulates cell survival, proliferation, differentiation and the maturation processes of mature megakaryocytes (MKs), and therefore the abundance of platelets in the blood^1^.

In a healthy organism, platelets and MKs take up TPO and degrade it, which leads to an inversely proportional occurrence of platelets and circulating TPO in the blood^4^. Moreover, Mpl is also a stem cell receptor, regulating haematopoietic stem cell (HSC) maintenance as well as expansion under inflammatory conditions or after transplantation^5–7^, suggesting a TPO-specific, demand-dependent activation of HSCs. However, the TPO stimulating effects on MKs are not yet clear. There is evidence that megakaryocytosis is inhibited by TPO-induced MK proliferation arrest and a promotion of terminal differentiation^8^. However, TPO signalling in MKs is dispensable for platelet production^9^.

Mutations or chronic stimulation can lead to a loss or to a gain of the Mpl-receptor function, which entails haematopoietic disorders such as congenital amegakaryocytic thrombocytopenia (CAMT) or myeloproliferative neoplasm (MPN), respectively. CAMT is a rare autosomal recessive bone marrow failure syndrome, which can evolve into aplastic anemia and leukaemia in early childhood^10^ and which results from mutations of the TPO receptor Mpl. In contrast, MPN is characterised by thrombocytosis and excess production of MKs and early progenitors^1^. Activating mutations of the Mpl receptor and TPO overexpression are known to contribute to the development of MPNs.

Mice with disturbed megakaryopoiesis/thrombopoiesis are important models to gain insights into the role of TPO stimulation of various cell stages and, eventually, to identify promising therapeutic targets for the aforementioned diseases. Mpl-deficient mice (*Mpl*^(−/−)^), which are used as a model for the CAMT phenotype, are characterised by thrombocytopenia accompanied by an overall loss of MKs and early progenitors. Restoring the Mpl-expression in bone marrow by means of gene therapy recovers thrombopoiesis, which indicates that the loss of HSCs in MPL-deficient mice is reversible^9, 11^. In contrast, mice with Mpl-deficiency in MKs and platelets only (*Mpl*^(*PF* 4*cre/PF* 4*cre*)^), exhibit a phenotype that is comparable to MPN^9^. For a detailed explanation of the mouse strains we refer to Ng et al.^9^. Alternatively, those clinical phenotypes can also be induced by a modified TPO expression (*TPO*^*Tg*^). While chronic excessive *in vivo* TPO stimulation (*overexpression*) leads to a MPN-like phenotype as in mice with Mpl-deficient MKs and platelets (*Mpl*^(*PF* 4*cre/PF* 4*cre*)^), TPO deficiency gives rise to a CAMT-like phenotype which is also exhibited by completely Mpl-deficient mouse strains (*Mpl*^(−/−)^)^9, 11^.

Several mathematical models have been developed to understand and quantify megakaryopoiesis and platelet production. Most of these models represent multi-compartment models focusing on the modulating effects of chemotherapy^12–14^ and radiation^15, 16^. Taking a different approach, Apostu and Mackey^17^ developed a rather simplistic model using a delay differential equation to understand oscillations observed in patients with cyclic thrombocytopenia. Although the delay component within this model is well suited to explain the cyclic behaviour of thrombocyte levels, the question remains whether the distinction in haematopoietic progenitors and MKs is sufficient to quantitatively explain how deregulation of the TPO-Mpl signalling affects the process of megakaryopoiesis and how the impaired but stable disease phenotypes result from TPO overexpression or knock out.

In order to answer this question, we take up the concept of this simplistic mathematical model and apply it to both the healthy situation and to two diseased phenotypes mimicking MPN. By evaluating the model’s ability to consistently describe those diverse outcomes as a result of deregulated TPO signalling, we obtain insight as to whether or not the simplistic distinction between progenitor and MK regulation is sufficient or not, or whether additional mechanisms need to be considered. For our analysis, we intentionally keep the model complexity as low as possible which allows us to focus on *essential* mechanisms and adhere to the fact that only relative (equilibrium) abundances of TPO, MKs, platelets and HSCs with respect to the control type are available.

## 1 Methods

### Experimental Data

The model analysis is based on experimental results from Ng et al.^9^, for which the authors comparatively studied different mouse strains with and without deregulated platelet production. For our analysis, we consider the following settings: (WT) wild-type mice with normally regulated haematopoiesis, (KO) conditional Mpl-knockout mice (*Mpl*^(*PF*4*cre/PF*4*cre*)^ with Mpl deficient megakaryocytes and platelets resulting in an MPN-like phenotype, and (OX) constitutive TPO over-expression *TPO*^*Tg*^ mice, also leading to an MPN-like phenotype. Our focus lies on TPO concentration, *T*, cell counts of primitive haematopoietic stem and progenitor cells, *H*, and megakaryocytes and platelets, *P*. Stable TPO levels and cell abundance for the mutated strains are given relative to WT controls. In particular, the following relative equilibrium abundances can be obtained from^9^: 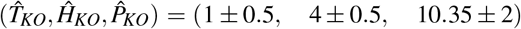 for the Mpl-KO genotype and 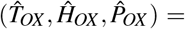 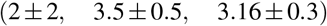 for *TPO*^*Tg*^ genotype exhibiting TPO overexpression. Of note, the standard errors are roughly estimated from the error bars of bar diagrams obtained from^9^ and should be taken here as guide to the range one has to reckon for the target values we want to reproduce with the mathematical model. For a detailed appraisal of the experimental conditions, please cf. the original publication^9^.

### Mathematical Model

Apostu and Mackey^17^ suggested a delay differential equation model of megakaryopoiesis/thrombopoiesis to account for the occasional occurrence of cyclic thrombocytopenia. We broadly follow this modelling approach in^17^, although we omit the delay part as no corresponding data on oscillations is available in the context of our research question. In brief, we describe megakaryopoiesis as a two step process, in which haematopoietic stem and progenitor cells (*H*) differentiate into megakaryocytes and platelets (cf. Figure 1). Herein we understand *P* as the lumped mass of Mpl-receptors carried on the surface of MKs and platelets. TPO (*T*) is constitutively expressed and regulates MK/platelet production as well as stem and progenitor cell proliferation via binding to its receptor, Mpl, expressed on the cells (depicted in blue in Figure 1).

**Figure 1.**
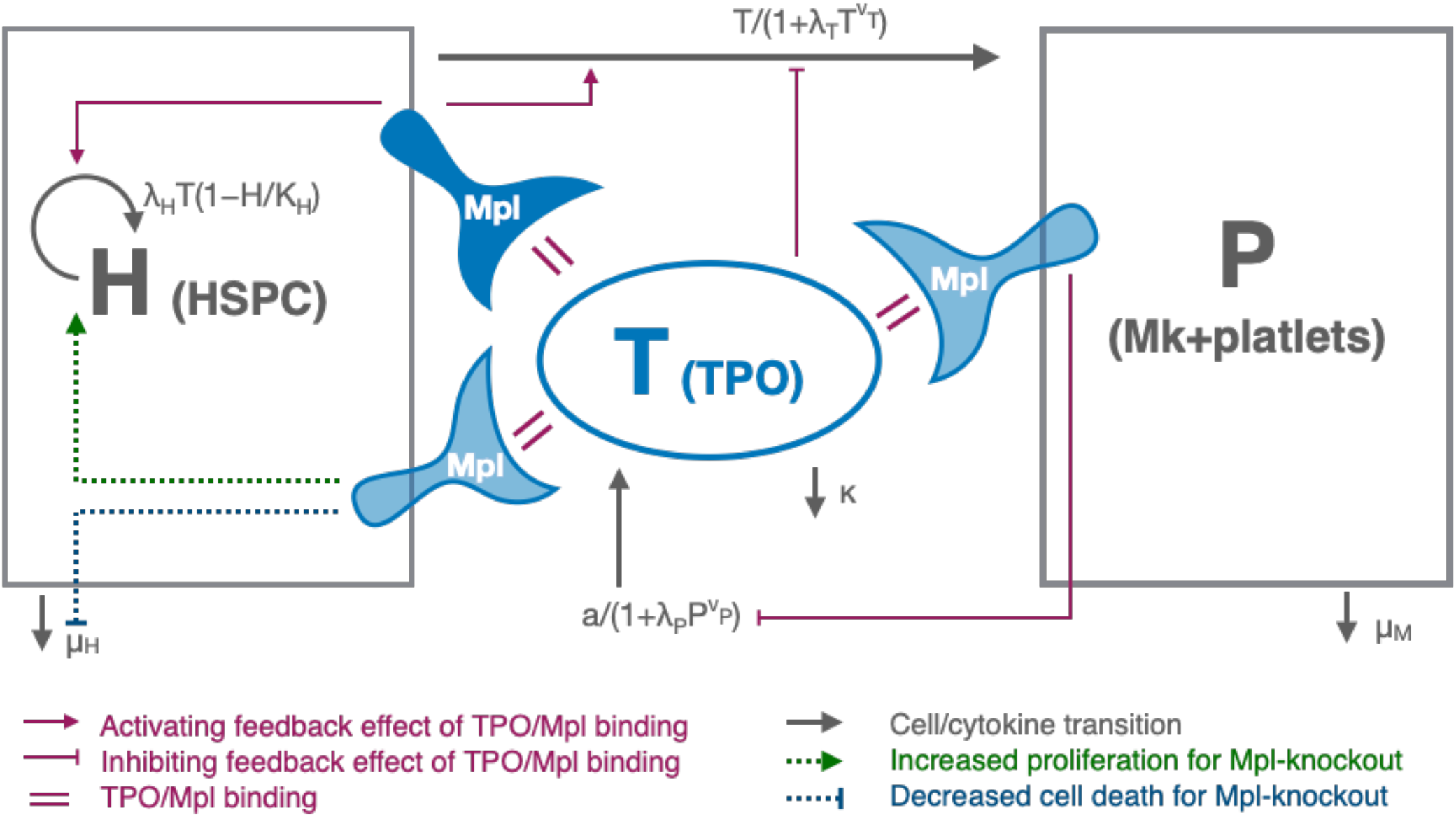
Scheme showing the regulations caused by Mpl-TPO ligation on haematopoietic stem and progenitor cells H and megakaryocytes and platelets P. Mpl receptors in light blue are affected by restricted Mpl KO, whereas the receptors depicted in dark blue are not affected by KO. Mpl-TPO ligation on P cells inhibits TPO production (parameter *λ*_*P*_ in eq. 3), and as a secondary effect inhibits differentiation of H cells / production of P cells (parameter *λ*_*T*_ in eqs. 5 and 6). The activating effects of Mpl-TPO ligation within the H-compartment (Mpl depicted in dark blue) on H-proliferation and Mk/platelet production, respectively, are not affected by restricted Mpl-KO (cf. eq. 5). We speculate, that a fraction of Mpl receptors, which are expressed on megakaryocytic progenitor cells within the H-compartment (depicted in light blue), are also deregulated and give rise to excess proliferation for the KO scenario.

We furthermore assume, that the relative abundances obtained from^9^ represent either stable equilibrium states or, in case of oscillations, represent averages over different cycles. In the latter case, the oscillatory amplitudes superimpose the observed variance.

To start with, the equilibrium value (or oscillatory average) of TPO concentration is described as an inverse function of the platelet abundance, which is given by

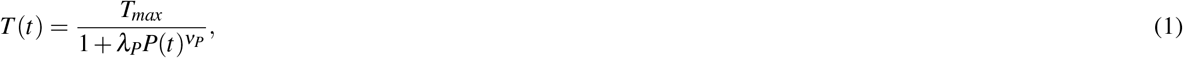

with *t* → ∞. Equation 1 can be found by evaluations of equilibrium dose-response/affinity assays. Typically, due to cooperativity of receptors expressed on single cells, more than one occupancies of the receptor-ligand compound can be observed, which gives rise to a Hill coefficient *ν*_*P*_ > 1. In addition, such an exponent may reflect a specific steric condition or multimeric receptor characteristics. Thus, the amount of produced receptor-ligand pairs proportional to the avidity parameter *λ*_*P*_ is not simply proportional to the receptor-expressing cell abundance, but rather to a power of the abundance.

From a dynamical perspective, eq. 1 results from the equilibrium condition 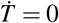 applied to differential equation describing the TPO-dynamics

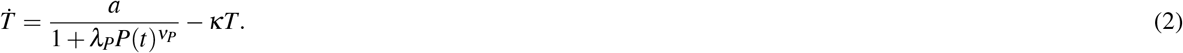

Thus, TPO concentration, *T*, has an upper bound 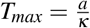 and eq. 1 then reflects the empirically validated inverse relation between TPO concentration and the mass of Mpl expressed by platelets and MKs. The inhibition of TPO-production emanating from the *P−*compartment is depicted in Figure 1. Parameter *λ*_*P*_ is a surrogate of the rate of degradation of TPO through interaction with Mpl expressed on platelets and MKs. In a simplistic interpretation, a Mpl-knockout mouse *Mpl*^(*PF* 4*cre/PF* 4*cre*)^ would be characterised by setting the rate of TPO degradation *λP* = 0, leading to a rise of TPO up to its maximum *T*_*max*_. However, as shown and discussed later in the results section, experimental data suggests that eq. 1 has to be modified by introducing a residual degradation parameter *ξ* leading to

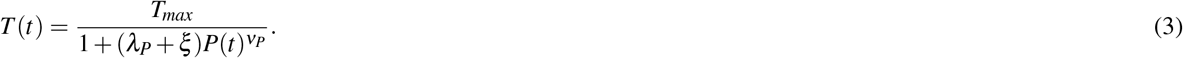

Since we here aim at reproducing relative abundances with respect to the “normal” homeostatic case, we set *T*_*max*_ := 1 for the upper limit of homeostatic TPO concentration.

The dynamics of the pool of haematopoietic stem cells and progenitor cells, capable to induce megakaryopoiesis with a subsequent differentiation of MKs to platelets, obeys the differential equation

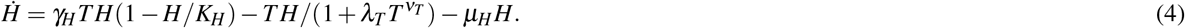

Since we deal with relative abundances, some of the estimated parameter values have to be understood relative to the arbitrarily chosen normalisations (without loss of generality). Only strain-dependent fold-changes are meaningful measures for those parameters. Specifically, the leading parameter of the middle term in eq. 4 has been set to unity. The proliferation (first) term is a logistic growth function with baseline proliferation rate *γ*_*H*_ and carrying capacity fixed to *K*_*H*_ = 100. Free estimation of proliferation rate *γ*_*H*_ and death rate *μ*_*H*_ suffices to fit the relative number of haematopoietic stem cells and progenitor cells, *H*, to the corresponding data. As depicted in Figure 1, proliferation of haematopoietic stem and progenitor cells is stimulated by an activating feedback effect of TPO-Mpl binding.

Differentiation of haematopoietic stem and progenitor cells (and therefore the production of MKs and platelets) is proportional to TPO concentration but limited by a Hill equation 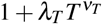 with two further free parameters *λ*_*T*_ and *ν*_*T*_. The latter term is a standard way of implementing density dependent kinetic rates. An increasing TPO concentration increasingly inhibits the production of MKs proportional to a receptor-ligand avidity parameter *λ*_*T*_ and proportional to a power of TPO concentration, 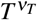, where *ν*_*T*_ is an occupancy parameter as explained for the case of *ν*_*P*_ in eq. 1 above. This TPO dependent inhibition, largely driven by TPO-Mpl ligation of cells in P, is depicted in Figure 1. Of note, exponents *ν*_*P*_ and *ν*_*T*_ are independent from normalisations.

Equilibrium condition 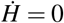 leads to the equilibrium value for the haematopoietic stem cells and progenitor cells

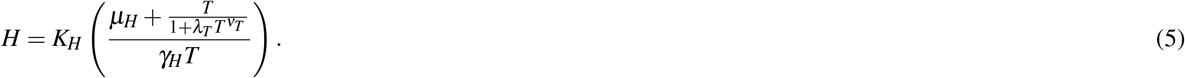

According to eq. 4, MKs are produced at rate 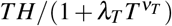. After a typical average delay time *τ*, MKs differentiate to platelets which are then produced at rate 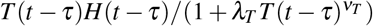 (cf.^17^). We neglect the delay *τ* to eventually yield

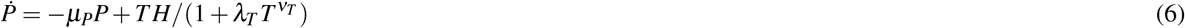

for the dynamics of *P*, interpreted as the “lumped mass” of Mpl-receptors carried on the surface of MKs and platelets, where *μ*_*P*_ is the relative death rate. In doing so, sustained oscillations are avoided, which we assume not to be relevant in the given context.

Thus, since we deal with relative equilibrium states only, the calculations needed within the optimisation procedure reduces to the computation of equilibrium values eqs. 3 and 5 for TPO, *T*, and haematopoietic stem and progenitors cells, *H*, respectively, whereas the solution for the platelet/MK dynamics is drawn from integrating differential eq. 6.

### Parameter optimisation using maximum likelihood

Based on the model given by eqs. 3, 5, and 6, we simulate three sets of strain-dependent time courses with the following strain-specific parameter settings:

WT: homeostasis with constant *T*_*max*_ = 1 and *K*_*H*_ = 100, and all other parameters freely varying.
KO: Mpl knockout with constant *T*_*max*_ = 1, *K*_*H*_ = 100, *λ*_*P*_ = 0, and *λ*_*T*_ = 0. Parameters *λ*_*P*_ and *λ*_*T*_ quantify the strengths of TPO-Mpl ligation (avidity), which is assumed to be zero for the Mpl knockout case while all other parameters can vary freely.
OX: TPO overexpression with constant *K*_*H*_ = 100, and all other parameters freely varying.

All other parameters than those defining the three strains are identical for all three settings. The relative equilibrium values (comparing KO and OX to WT) are obtained by solving equations 3, 5, and 6 for KO and OX to obtain their homeostatic equilibrium and relating those values to the corresponding solution for the WT condition. During the optimisation process, all except the strain-specific parameters are varied simultaneously for the three scenarios. From the resulting solutions we obtain relative equilibrium values which are then compared to the observed relative abundances 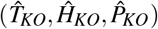 and 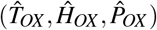, respectively. To this end we apply a maximum likelihood procedure, thus allowing for an estimation of the *optimal* parameter values which minimise the differences between model solution and observed abundances. Deviations from these strain-dependent parameter settings are discussed in the Results section.

A direct evaluation of the equilibrium states of equations 3, 5, and 6 leads to a set of implicit equations for the parameters, for which an analytic solution appears not to be feasible. Therefore we employ a numerical approach to solve the equations and identify the steady state values of each simulation with different parameter values. Starting on the initial condition *P*(0) as well as the given parameter values, the solutions of eqs. 3, 5, and 6 exhibit transient curves that converge towards the equilibrium values, often in form of damped oscillations. A time series of an arbitrary length of 10,000 time steps is computed, evaluated at each time step. Although the curves quickly converge to equilibrium roughly within 10 to 100 time steps, the first half of the simulation is skipped and the deviations of the remaining curves from the experimentally observed values are evaluated and fed into the maximum likelihood routine. For an illustration of this procedure, cf. Figure 2.

**Figure 2.**
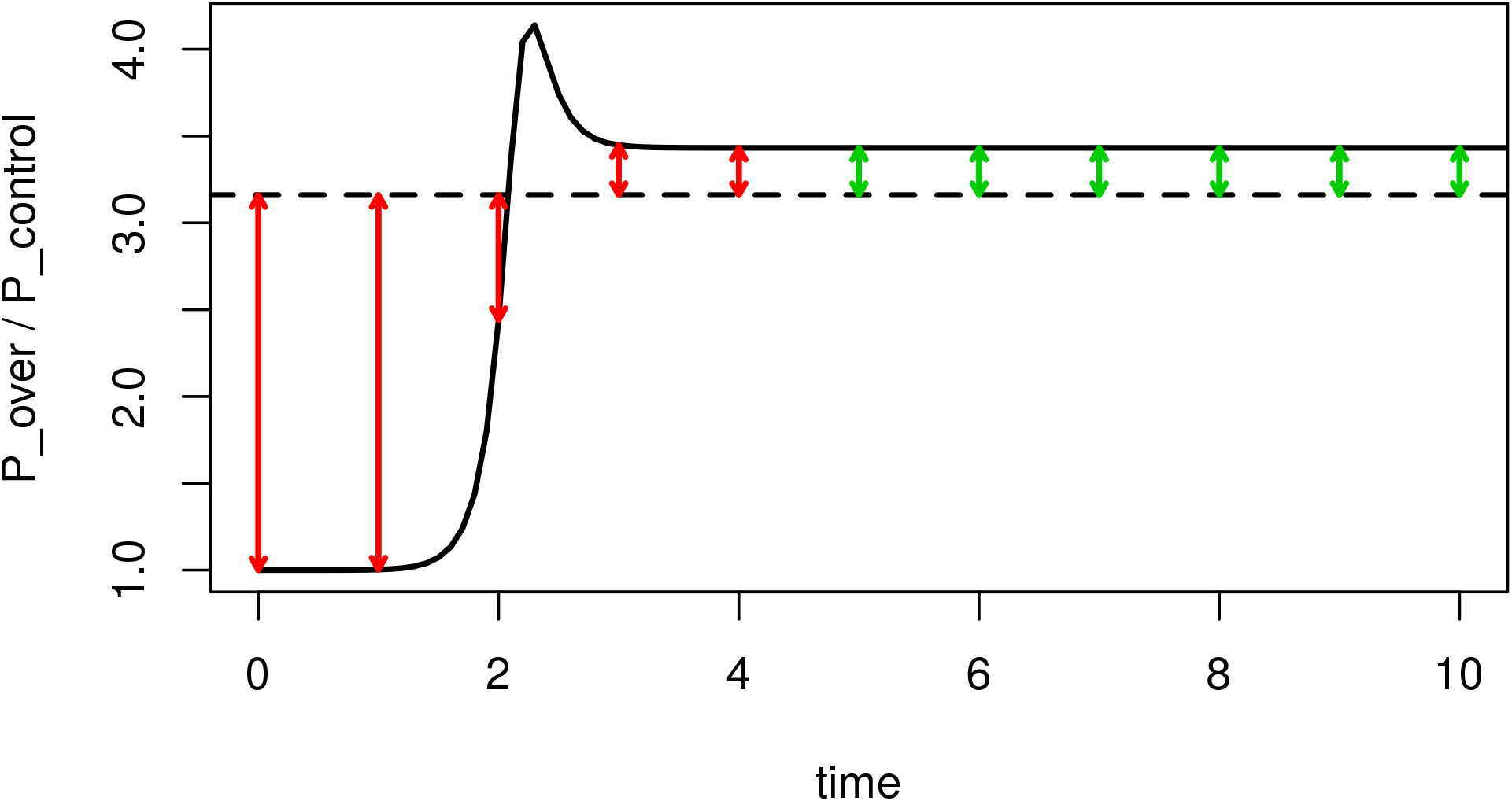
Illustration of the fitting procedure. The dashed line represents the reference value of relative platelet/MK abundance 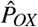 for the case of TPO overexpression. The bold curve shows a model prediction exhibiting a short damped oscillation in the beginning with subsequent convergence towards a steady state. Thereby, suboptimal values for the kinetic parameters have been used, thus leaving deviations with respect to the reference curve, which are fed into the maximum likelihood routine in order to be minimized. Minimizing all residuals starting from *t* = 0 (red and green arrows) leaves a short transient oscillation, whereas using only deviations at later time points (green arrows) for minimization allows for a more sustained oscillation.

In order to assure that the actual time length used for the model adaptation does not severely influence the parameter estimation, we fitted the model for different lengths of the simulated time series. Several test cases with varying simulation times did only marginally affect those estimates. In particular, we observed that the duration of the fitting interval (see Figure 2) influences the magnitude and duration of transient oscillations before the system converges into the steady state solution. However, this effect does neither influence the model selection procedure nor the asymptotic steady states.

## 2 Results

### Inability to consistently describe TPO-Mpl knockout and overexpression

We study whether a two compartment model of megakaryopoiesis is quantitatively sufficient to describe both alterations that result from knockout (KO) of the TPO-Mpl signalling in stem cells and megakaryocytic progenitors or from its continuous overexpression (OX). In terms of the mathematical formulation (see section 1, Mathematical Model), both TPO overexpression as well as Mpl-knockout translate into distinct modifications of certain parameter values. As such, we assume that the rate of TPO degradation by platelets and MKs (compartment *P* in model equation 3) differs between control mice and the *Mpl*^(*PF*4*cre/PF*4*cre*)^ strain. Specifically, and assuming an efficient knockout, the rate of TPO degradation should drop to zero in Mpl-deficient mice. Other kinetic parameters are expected to be identical for the different mouse strains. Under those assumptions, a sufficient model should be able to reproduce the relative abundances of TPO concentration *and* cell numbers observed for the different mouse strains. If the model fails to consistently describe the data, we take this as an indication that further opaque mechanisms influence the quantitative outcomes.

Applying the optimisation routine for the strain-dependent parametrisation as given in section Methods (Parameter optimisation using maximum likelihood), we obtain a perfect fit for the relative platelet abundances (compartment *P* in model eq. 6). Varying the target values 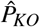 and 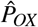 within the range of the observed biological variability (cf. section Methods, Experimental Data) likewise leads to perfect fits for the relative platelet counts. In other words, our model with the suggested strain-dependent parametrisation is flexible enough to reproduce platelet counts for both the KO and the OX scenario. Of note, *λ*_*T*_ = 0 (indicating a complete KO, thus a removal of the inhibitory effect of TPO on MK/platelet production) for the *Mpl*^(*PF* 4*cre/PF* 4*cre*)^ mouse line is a necessary condition to explain the observed platelet data. In addition, a residual inhibition of TPO concentration through platelets expressed by a non-negligible value for parameter *ξ* is indispensable to ensure an acceptable quality of the fit (see Table 1 for the parameter estimates).

**Table 1.** List of parameter estimates. Six parameters have either been fixed (*K*_*H*_ = 100) for all genotypes or have been estimated independently from the genotype (5 bottom rows). Four parameters (4 top rows) and an additional fold-change of *γ*_*H*_ have been stratified for the genotypes. For details cf. text.

| | control | $TPO^{Tg}$ | $Mpl^{(PF4cre/PF4cre)}$ |
| --- | --- | --- | --- |
| $T_{max}$ | 1 (fixed) | 2.90 | 1 (fixed) |
| $\lambda_P$ | 3.08 | | 0 (fixed) |
| $\lambda_T$ | 10.62 | | 0 (fixed) |
| $\gamma_H$ | 11619.91 | | $2.5 \cdot 11619.91$ |
| $K_H$ | 100.00 (fixed) | | |
| $v_T$ | 1.58 | | |
| $\mu_H$ | 2898.33 | | |
| $\mu_P$ | 10.53 | | |
| $v_P$ | 0.43 | | |
| $\xi$ | 1.78 | | |

In contrast, the predicted relative counts of haematopoietic stem cells and progenitor cells (compartment *H* in model eq. 5) are below the observed values for the Mpl-KO case (*H* = 3.32 vs 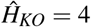) and above the corresponding observation for the case of TPO overexpression (*H* = 4.15 vs 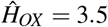). A similar pattern can be observed for the TPO concentration, *T*. The predicted relative TPO concentration is above the observation for the Mpl-KO case (*T* = 1.39 vs 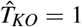) and below observation for the case of TPO overexpression (*T* = 1.62 vs 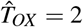). These results entail a missing link in the structure of the model or the parametrisation. We conclude, that the model in the current form is not flexible enough to consistently reproduce relative TPO concentration, relative platelet counts and relative stem cell abundances.

### Strain-independent model adaptations cannot improve quantitative fits

We further investigate whether missing links between TPO concentration and the cell compartments can account for the observed differences between model and data. We report about several attempts to improve the fitting quality by adding strain-independent terms to the dynamic equations. For example, adding the two kinetic parameters *η* and *T*_*min*_ to the equilibrium TPO value in form of

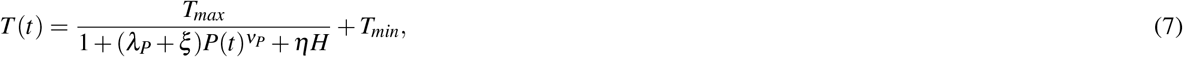

did not improve the quantitative model fits. In particular, the assumption that TPO has a minimum equilibrium value, *T*_*min*_, independent of any control through the number of available MK/platelets leads to an estimate of *T*_*min*_ ≃ 0 and does not improve the likelihood of the model. The same holds for *η*, a parameter that takes into account a control of TPO concentration by haematopoietic stem cell abundance *H*.

We further tested whether a mechanism that accounts for the chalon-mediated inhibition of MK growth resulting from destructed platelets^18^, can be incorporated into our model to enhance the fitting quality. Technically, this can be achieved by adding a corresponding influence of the platelet abundance *P* in three different ways, not necessarily simultaneously (see bold fonts):

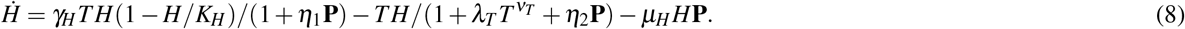

Of note, *η*_2_**P** has to appear in the corresponding transfer rate for the platelet production, too. None of the three terms shows an effect. Substituting *η*_1_**T** for *η*_1_**P** likewise has no impact. We refrain from reporting further unsuccessful attempts to modify the model in a strain-independent manner. We conclude, that adding or altering terms of the model does not improve the outcome, even though the alterations come with additional parameters and increase the degrees of freedom.

### Strain-dependent model adaptations explain quantitative differences and hint toward previously unconsidered Mpl knockout effects

Based on the above results we argue that there must be an additional strain-dependent parameter which has not been considered so far. In order to explore such mechanisms we release our assumptions and study the case, in which the Mpl-KO, but not the OX, also affects stem cell proliferation. We depict this additional mechanism by the dashed lines in Figure 1, explaining the excess proliferation within the H-compartment for the Mpl-KO case.

Technically, we introduce a factor accounting for KO-specific excess proliferation *α* of the haematopoietic stem and progenitor cells *H*. Keeping this factor at *α* = 1 for the WT and the OX scenario, we find that *α*_*KO*_ = 2.5 for the KO scenario results in a perfect fit of the model to the observed relative equilibrium values. Moreover, if the target values 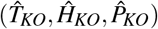 and 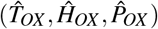 are varied over the range of their observed variability (cf. section 1, Experimental Data, and ref.^9^ for the standard errors of the 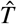−, 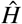− and 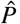− values), the corresponding estimated equilibrium states exactly match with these reference equilibria. Thus, the introduction of an additional strain-dependent parameter renders the model suitable to explain the entire observed biological variability of the relative homeostatic values. Although a strain-dependent increase in the proliferation rate of the stem cells consistently resembles the observed phenotypes, we are left with the question how such a more than 2-fold increased proliferation rate for the Mpl-knockout can be explained.

Within our model, proliferation and death rate are intrinsically entangled. Therefore, the corresponding parameters *γ*_*H*_ and *μ*_*H*_ cannot be estimated independently. As a consequence of this technical ambiguity, an *α*−fold increase of the proliferation rate *γ* leads to the same fit as a 1*/α* reduction of the death rate *μ*_*H*_. For the given model structure we cannot reveal whether the excess increase of HSCs *α*_*KO*_ = 2.5 in the *Mpl*^(*PF*4*cre/PF*4*cre*)^ mouse strain derives from an increase of the proliferation rate or a decrease of the death rate. The complete set of parameter estimates is listed in table 1 for the case of estimating the fold-change of *γ*_*H*_ for the *Mpl*^(*PF*4*cre/PF*4*cre*)^ mouse line. Removing the factor of *α*_*KO*_ = 2.5 and instead dividing *μ*_*H*_ by 2.5 for this mouse strain reflects the competing scenario.

We conclude that an excess proliferation (or decreased cell death) of haematopoietic stem and progenitor cells appears as a side effect of the Mpl-KO, thereby further altering the steady state equilibrium values.

## 3 Discussion

The presented three-dimensional model for megakaryopoiesis (eqs. 3, 5 and 6) accounts for the population dynamics of TPO *T*, haematopoietic stem and progenitors cells *H*, and platelets/MKs *P*, respectively. In its baseline version it has a total of 9 free kinetic parameters, such as proliferation and death rates. If this model sufficiently represents the major underlying regulations, there should be a set of optimal parameters such that the resulting model simulation accounts for the WT control scenario as well as the modified mouse strains in which TPO-Mpl signalling is altered. In particular, we chose two different mouse strains, (*Mpl*^(*PF*4*cre/PF*4*cre*)^ and *TPO*^*Tg*^), both of which mimic the pathological phenotype MPN exhibiting a perturbed homeostasis. We would expect that the homeostatic, steady state values of TPO concentration, stem cell counts and platelet counts obtained from the model obey the same relative abundances as observed for the experimental situations^9^. These sets of optimal parameter values have been estimated in an indirect way by fitting the model to the relative abundance with respect to the homeostatic values of the WT control.

In order to reproduce the observed abundances for the case of TPO OX (reflecting the transgenic mouse strain *TPO*^*Tg*^), we expected, by virtue of biological plausibility, that it suffices to adapt only one parameter, namely *T*_*max*_ in eq. 3, to an appropriate value. This conjecture could be confirmed since the other 8 parameters are invariant upon switching from the healthy controls to the *TPO*^*Tg*^ mouse line. Thus, TPO OX can be accurately explained by an increased maximum TPO level.

In an analogous fashion we investigated whether a knockout of TPO receptors expressed on platelets and MKs (mouse line *Mpl*^(*PF*4*cre/PF*4*cre*)^) can be explained by eliminating the TPO-Mpl signalling (i.e. setting reaction kinetic parameters corresponding to the Mpl affinity for its ligand TPO to zero). However, the identification of these parameters was not as straightforward as for the TPO overexpression case. Setting *λ*_*P*_ = 0 in eq. 3 for the Mpl-KO case turned out to be necessary but not sufficient to adequately tune the model to the desired behaviour. Additionally setting *λ*_*T*_ = 0 in eqs. 5 and 6 substantially improves the fit but is still not fully satisfactory. Parameter *λ*_*T*_ appears in the conversion rate of HSCs to platelets and, therefore, quantifies transient MK-production. Specifically, the term 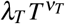 quantifies internalisation and degradation of TPO by MKs. Thus, it appears to be biologically plausible to assume *λ*_*T*_ = 0 for the Mpl-KO case.

The inability of the model setup to consistently explain different phenotypes only as the consequence of disabled TPO-Mpl signalling points to the main conclusion of our study. We reason that further mechanisms within the Mpl knockout strain have an additional impact on renewal or/and survival of haematopoietic stem cells and progenitor cells. Integrating this assumption into the model framework by accounting for an excess proliferation factor *α*_*KO*_ = 2.5 (or a corresponding 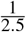− fold reduction of the death rate) leads to a perfect correspondence between model predictions and experimental observations.

We may speculate about the biological interpretations of this opaque mechanism. To this end, we recall that the tripartition of megakaryopoiesis into HSCs, MKs, and platelets is rather simplistic. Furthermore, we have lumped the transient MK state and platelet abundance into one compartment. Therefore, decreased removal rates of HSCs after Mpl-KO could also reflect an increased removal of early MK precursors due to MK-TPO interaction. We further raise the question whether such a differential response may also result from altered Mpl expression among the haematopoietic stem cells and progenitor cells themselves. Although it has been reported that Mpl-knockout in the *Mpl*^(*PF*4*cre/PF*4*cre*)^ mouse line is stringently restricted to platelets and MKs, relaxing this assumption would represent another possible and quantitative explanation of the observed phenotypes.

To conclude, we supply strong evidence that the well-known excess production of MKs and early progenitors^1^ in MPN is either directly contingent on Mpl expression on platelets and MKs or, alternatively, that the knockout of Mpl in *Mpl*^(*PF*4*cre/PF*4*cre*)^ mice is not as precisely restricted to MKs and platelets. Our results contribute to the ongoing discussion on whether the functional distinction between MKs and their progenitors occurs on a stepwise manner or whether it should be understood as a gradual transition^8^. Mathematical models represent important tools to test the quantitative correctness of conceptual models and to propose additional regulatory mechanisms thereby guiding further experimental investigations.

## Acknowledgements

This work was supported by the German Federal Ministry of Education and Research (www.bmbf.de/en/), Grant number 031A315 “MessAge” to I.G.

## Author contributions statement

A.G. conceived the work. H.H.D. conceptualised the mathematical model and drafted the manuscript. All authors reviewed and revised the mathematical model. I.G. supervised the work. All authors reviewed and revised the manuscript.

